# Adaptive FDR thresholding of Fourier shell correlation for resolution estimation of cryo-EM maps

**DOI:** 10.1101/2020.03.13.990705

**Authors:** Maximilian Beckers, Carsten Sachse

**Affiliations:** European Molecular Biology Laboratory (EMBL), Structural and Computational Biology Unit, Meyerhofstraße 1, 69117 Heidelberg, Germany; Candidate for Joint PhD degree from EMBL and Heidelberg University, Faculty of Biosciences; Ernst-Ruska Centre for Microscopy and Spectroscopy with Electrons (ER-C-3/Structural Biology), Forschungszentrum Jülich, 52425 Jülich, Germany; JuStruct: Jülich Center for Structural Biology, Forschungszentrum Jülich, 52425 Jülich, Germany

**Keywords:** Cryo-EM, Fourier shell correlation, resolution criteria, permutation sampling, false discovery rate

## Abstract

Fourier shell correlation (FSC) has become a standard quantity for resolution estimation in electron cryo-microscopy. However, the resolution determination step is still subjective and not fully automated as it involves a series of map interventions before FSC computation and includes the selection of a common threshold. Here, we apply the statistical methods of permutation sampling and false discovery rate (FDR) control to the resolution-dependent correlation measure. The approach allows fully automated and mask-free resolution determination based on adaptive thresholding of FSC curves. We demonstrate the applicability for global, local and directional resolution estimation and show that the developed criterion termed FDR-FSC gives realistic resolution estimates based on a statistical significance criterion while eliminating the need of any map manipulations. The algorithms are implemented in a user-friendly GUI based software tool termed SPoC (https://github.com/MaximilianBeckers/SPOC).

## 1. Introduction

Electron cryo-microscopy (cryo-EM) is becoming established as one of the methods of choice for macromolecular structure determination. The technique has undergone major improvements in hardware and software, allowing to determine 3D structures routinely at close-to-atomic resolution (Kühlbrandt *et al.*, 2014; Bartesaghi *et al.*, 2015; Weis *et al.*, 2019). These resolutions provide the landmark features for atomic model building and the resulting atomic coordinates rationalize biological mechanism and function. At the core of all these developments are improvements in the resolvability of molecular detail that can be achieved. Therefore, the resolution is a reported value that makes the data comparable with other structural biology methods, and more importantly, gives guidance how confidently we can interpret the density of a given feature in 3D space. Fourier shell correlation (FSC) curves are the standard metric for resolution estimation of cryo-EM maps. FSCs were originally introduced to the field of cryo-electron microscopy (Harauz & Van Heel, 1986) and more recently were also applied to related fields such as super-resolution microscopy (Nieuwenhuizen *et al.*, 2013; Banterle *et al.*, 2013).

The FSC measures the correlation between Fourier coefficients within every resolution shell of two independently determined half-maps. A typical curve shows high correlations at low resolutions until it drops to zero at higher resolutions when noise starts to dominate the signal. In order to report a resolution value of structures based on FSC curves, a threshold value needs to be selected. A fixed threshold of 0.143 has been proposed (Rosenthal & Henderson, 2003), which is widely used for resolutions better than 10 Å when map features can be used to validate the obtained resolution. For lower resolutions, a more conservative 0.5 threshold has typically been used (Spahn *et al.*, 2004). The 0.5 threshold is also favored when local FSCs within small windows are computed and local resolution is estimated (Cardone *et al.*, 2013). Nevertheless, fixed value thresholds ignore the effective number of Fourier coefficients in the respective resolution shell (Van Heel & Schatz, 2005). This effect becomes particularly relevant at local resolution estimation with small window sizes (Cardone *et al.*, 2013) and for 3D FSC with small number of Fourier pixels (Zi Tan *et al.*, 2017). Alternatively, other criteria like the 2*σ* and 3*σ* (Saxton & Baumeister, 1982; Orlova *et al.*, 1997) as well as the half-bit criterion have been proposed that compensate for these effects (Van Heel & Schatz, 2005). More generally, applying thresholds such as the 0.143/ 0.5 or half-bit criterion are to be interpreted as identifying the highest resolution shell that still contains a minimum of interpretable signal.

In order to get correct estimates of resolution-dependent information measures of the molecular density within a 3D reconstruction, solvent noise needs to be removed from the volume, as big parts of the reconstructed volume correspond to noise only and thus bias the resolution towards lower values. Therefore, mask application is critical, yet poses the danger of introducing artificial correlations. This effect has been proposed to be corrected by mask deconvolution (Chen *et al.*, 2013). In practice, this approach still involves testing of several masks until the user reaches an agreement between the displayed molecular features and the estimated resolution. Although the calculation of corrected FSC curves considering the molecular mass have been proposed to circumvent this problem (Sindelar & Grigorieff, 2012), such an approach is also used in combination with a defined threshold criterion.

Alternatively, *σ*-thresholds were proposed in order to provide cutoffs when the FSC exceeds the random correlations of pure noise (Saxton & Baumeister, 1982; Van Heel & Schatz, 2005). However, statistics based on simple *σ*-thresholds suffer from several drawbacks as linking *σ*-levels to probabilities requires strict assumptions about the underlying probability distributions, which are usually not known. In order to minimize subjectivity and increase robustness of FSC computations, we developed a new approach of adaptive thresholding for resolution determination using a combination of permutation sampling and *p*-value correction by false discovery rate control. We show that the procedure is able to detect signal with high precision over a wide range of noise levels that make masking of half-maps obsolete. We validate the approach by comparing a large batch of structures with reported resolution values of the EM databank and further extend the approach successfully to challenging cases of local and directional resolution determination.

## 2. Results

### Resolution estimation using statistical thresholding of the FSC

In order to circumvent principal and practical problems of thresholding FSC curves, we developed a procedure for identifying the highest resolution shell of interpretable signal based on parameter-free permutation sampling and subsequent statistical inference. Permutation sampling tests are statistical procedures for estimating the null-hypothesis from the data itself and thus do not require any assumptions about the underlying distributions (Lehmann & Romano, 2005). Permutation testing of FSC coefficients for each resolution shell is straightforward: under the null-hypothesis the two resolution shells from the half-maps are independent, we can thus generate new samples by permutation, i.e. scrambling the Fourier coefficients of the second resolution shell (**Figure 1a**). As a result, we compute a large series of noise FSCs. Effectively, this permutation allows sampling of the null distribution and as a result FSC-values for each resolution shell can be conveniently transformed into *p*-values. In order to account for this multiple testing problem, *p*-values are then corrected by means of false discovery rate (FDR) (Benjamini & Hochberg, 1995) control and thresholded at 1 %, as previously applied to cryo-EM map thresholding (Beckers *et al.*, 2019). We term this approach the FDR-FSC method of resolution determination.

**Figure 1.**
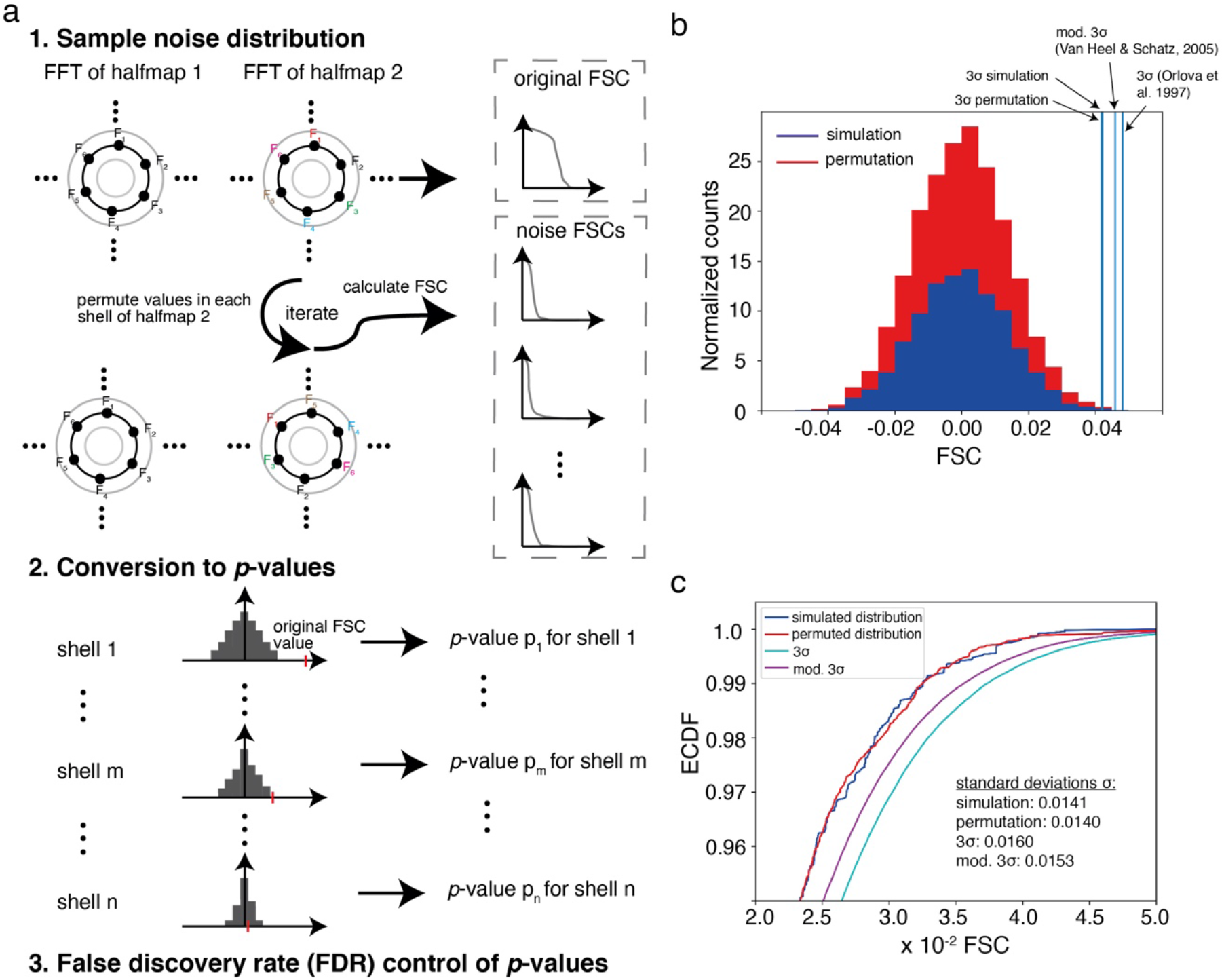
Resolution estimation by permutation-based FDR-FSC. (a) Samples of random noise FSCs are generated by permuting Fourier coefficients in half-map 2, creating newly paired Fourier coefficients (1.). The resulting random noise FSC distribution for each resolution shell is used to estimate the significance, i.e. a *p-*value for the original FSC value of each resolution shell (2.). The resulting *p*-values per resolution shell are subjected to false discovery rate (FDR) control (3.). (b) Stacked histogram of FSC values of independent half-maps from the half-Nyqist resolution shell (1/4 pixel) together with the respective 3*σ* cutoffs and (c) comparison of the empirical cumulative distribution functions (ECDFs) in a zoomed view. The true distribution sampled by multiple simulations of two noise maps and subsequent FSC calculation is shown (blue). The distribution obtained with our permutation approach is shown (red). ECDFs from normal distributions with standard deviations from the *σ*-threshold criteria are shown in blue (modified 3*σ* criterion) and cyan (original 3*σ* criterion). The true and the permutation-based distributions follow each other closely, especially at the tail of the distributions. Moreover, the modified 3*σ* is closer to the simulated distribution than the original 3*σ*.

In order to test whether the permutation approach captures the principal distribution of FSC coefficients, we simulated two pure noise half-maps with an average of 0 and a standard deviation of 1 and applied the permutation sampling as described. To assess the true distribution of FSC coefficients, we simulated 5,000 noise half-maps (again with an average of 0 and a standard deviation of 1) and calculated FSC curves for each pair of 5,000 half-maps. Comparison of the simulated with the permuted histograms generated from the FSC values of the half-Nyquist resolution shell (1/4 pixel) and the corresponding empirical cumulative distribution functions (ECDF) show that the distributions from the permutation approach and the simulation follow each other closely (**Figure 1b**), in particular at the tail of the distributions (**Figure 1c**). However, the comparison within the half-Nyquist shell shows that simple standard deviations computed from non-permuted FSC distributions as they are used by the proposed 3*σ* (Orlova *et al.*, 1997) and modified 3*σ* criteria (Van Heel & Schatz, 2005) for resolution estimation can give rise to overestimates in comparison with permuted distributions. Next, we wanted to assess the performance over all resolution shells and plotted the right-sided 10, 5.0, 1.0 and 0.15 % cutoff values (percentiles) for each resolution shell. The percentiles of the distributions from the permutation and the simulation follow each other closely (**Supplementary Figure 1a**). Together, the permutation approach within each resolution shell is able to accurately represent the FSC distribution under the null hypothesis and the standard deviations computed from the permuted distributions are more robust than simple standard deviations as they are used for traditional 3*σ* criteria.

When symmetry is applied during the image reconstruction, additional dependencies between half-maps are introduced. These effects on the FSC computation and cutoff determination have been discussed previously and can be compensated by the reduction of the sample size (Van Heel & Schatz, 2005). Therefore, in the permutation approach we took this effect into account and reduced the sample size from each resolution shell by the factor of available symmetry-related asymmetric volume units, i.e. D4 symmetry has 8 asymmetric volume units and reduces the effective sampling to 1/8 or 12.5 % of the initial sample size. In order verify the approach, we tested permutation-based sampling of two symmetry-imposed half-maps in the presence and absence of the sample size correction factor. In analogy to the test on the permutation approach above, we compared the permuted distributions with the ones obtained 5,000 noise symmetrized half-maps by plotting the 10, 5.0, 1.0 and 0.15 percentiles as a function of spatial frequency. In the absence of any sampling correction, the permuted and simulated percentiles do not overlap whereas as in the presence of a D4 and D7 symmetry correction factor, the permuted and simulated percentile lines approach each other more closely (**Supplementary Figure 1b** **and** **c**). Similar effects of introducing additional dependencies between half-maps occur during masking, which most image reconstruction algorithms make use of already by reconstructing density within a sphere. This has been discussed in (Van Heel & Schatz, 2005) as the filling degree of the reconstructed object in the volume. In order to compensate for such additional effects, we estimated the effective sample sizes from spherical masking. We minimized the deviation between the true distribution and the sample-size corrected distribution by testing systematically effective sample size factors between 0 and 1 and computing the Kolmogrow-Smirnow distance of the two distributions (Massey, 1951) (see Methods for more details). Using this approach, we found that a soft circular mask of volume size diameter reduces the effective sampling to 70% of the initial sample size (**Supplementary Figure 2a)** and a Hann window of half the volume dimension to 23 % of the initial sample size (**Supplementary Figure 2b**). The utility of the sample size compensation is demonstrated by the comparison of permuted and simulated percentiles using a series of different window sizes: the resulting percentile lines follow each other closely only in the presence of sample size correction (**Supplementary Figures 3** **and** **4**). In turn, these tests also show that any map manipulations e.g. by user-defined masking will affect the FDR-FSC measurements and should be avoided. In conclusion, operations that introduce additional dependencies between half-maps such as volume symmetrization and masking require compensation by using a sample size correction factor.

Based on the proposed FDR-FSC approach, the resolution estimate will now be assigned to the highest spatial frequency that contains significant signal at 1% FDR (from now on referred to as FDR-FSC). For conventional FSC computation, the half-maps are masked by the user, whereas for the here proposed FDR-FSC approach the untreated half-maps are used for FSC determination. The resolution value will be assigned to the resolution shell that crosses the significance threshold for the first time (**Figure 2a**). Comparison of the conventional 0.143 cutoff values thresholds for a *γ*-secretase map (Bai *et al.*, 2015) (EMD-3061) at 3.4 Å obtained by masking with the 3.4 Å FDR-FSC value obtained without any masking indicates a very similar resolution. For a series of EM databank (EMDB) entries (EMD2677, EMD0587, EMD4589, EMD0043, EMD3061, EMD0415, EMD6287, EMD8908), we used the raw half-maps for FSC computation and we assessed the influence of the chosen FDR threshold on the resolution measurement. For FDR cutoff values between 0.1 and 10%, the estimated resolution remains constant in most cases. As the FDR-FSC method is robust towards error thresholds between 0.1 and 10% (**Supplementary Figure 5a**), we continue to use the 1% FDR threshold for the FDR-FSC resolution estimation. To test the sensitivity towards noise, we reduced the window size of the volume of the same *γ*-secretase map in steps of 20 voxels, only removing solvent noise. It is evident that FSC curves move towards the Nyquist frequency as more noise is removed (**Figure 2b**). While the 0.143 FSC threshold decreases from 4.1 Å to 3.7 Å (**Figure 2c**), the adaptive 1% FDR threshold practically remains constant and only fluctuates at the second decimal digit. To further validate the robustness towards different noise levels, we simulated a map of *β*-galactosidase (PDB ID 5a1a (Bartesaghi *et al.*, 2015)) at 2.5 Å resolution and added different amounts of white Gaussian noise. Similar to our results from above we found that the resolutions at the 0.143 FSC criterion are very sensitive to the added noise and change already beyond 1.1 standard deviations (**Supplementary Figure 5b**), whereas the resolution determined using 1% FDR-FSC threshold remains constant at noise levels up to 4.0 standard deviations when high-resolution noise entirely dominates visibility of the structural features (**Supplementary Figure 5c**). These experiments show that the 1% FSC-FDR threshold gives realistic resolution estimates in agreement with user reported values from the EMDB and is more robust than the commonly used 0.143 criterion at much higher levels of noise present in the structure.

**Figure 2.**
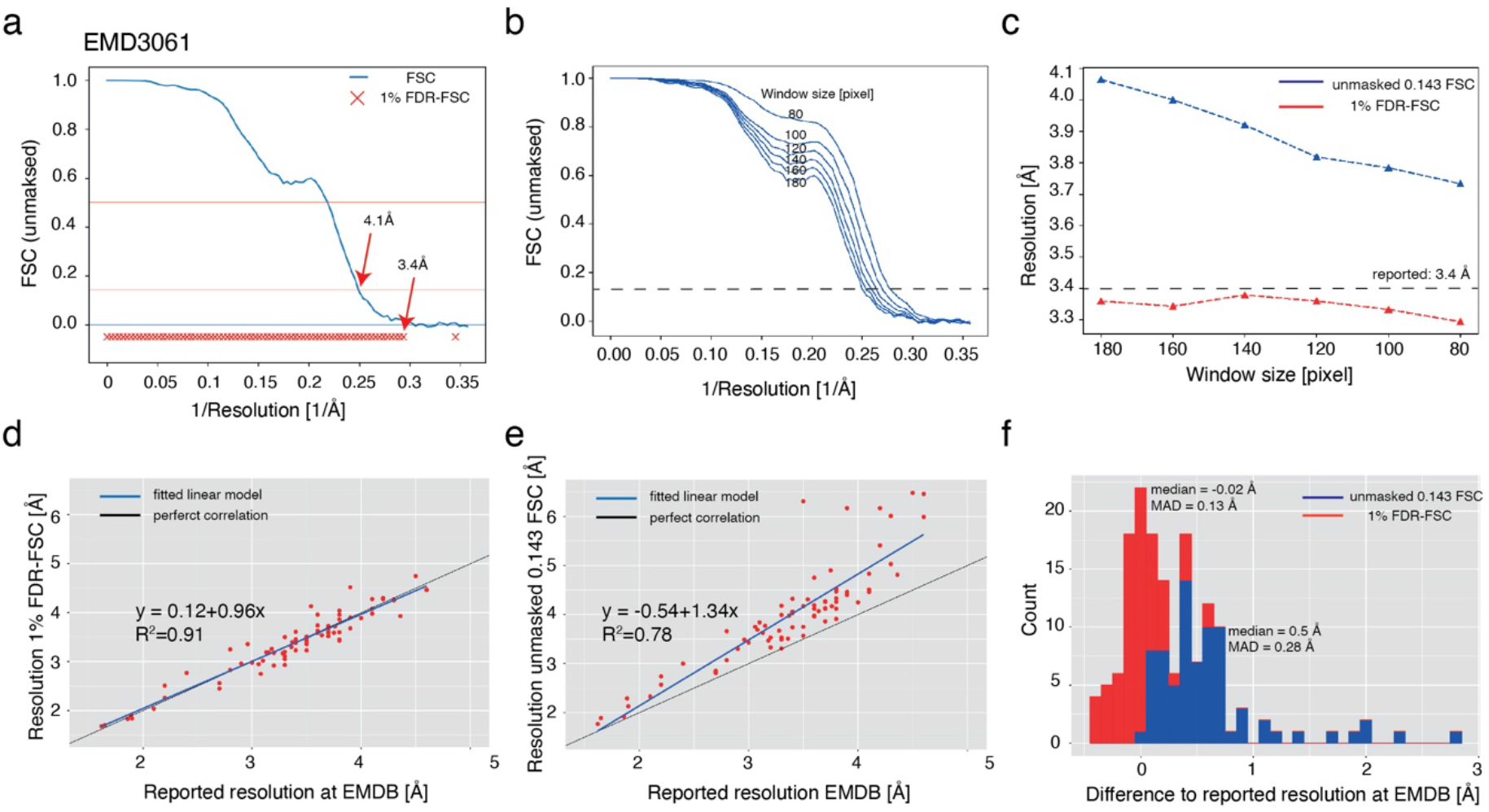
FDR-FSC results: effect of noise removal on FDR-FSC and benchmarking resolution estimation using 77 maps from the EMDB. (a) Example of a Fourier shell correlation (FSC) curve for EMD3061 computed from two untreated half-maps. Resolutions shells with significant correlations beyond random fluctuations at 1% FDR-FSC are marked with red crosses. (b) Effect of noise removal on FSC curve by decreasing window sizes in steps of 20 voxels and thereby incrementally excluding solvent noise. (c) Effect of noise removal on resolution estimates at 0.143 FSC (blue) compared with 1% FDR-FSC (red). (d) Scatter plot of reported resolutions against resolutions determined by the 1% FDR-FSC method. Fitted line (R^2^=0.91) is shown in blue. For comparison, perfect correlation is shown in black. (e) Scatter plot of reported resolutions against unmasked 0.143 FSC values. (f) Stacked histogram of resolution differences of the 1% FDR-FSC and the unmasked 0.143 cutoff with respect to the reported resolutions (blue and red).

### Performance comparison of FDR-FSC with common fixed threshold FSC resolution determination

In order to benchmark of the proposed algorithm, we used 77 reference raw half-maps from the EMDB, determined the resolution using the FDR-FSC approach in a fully automated fashion and compared them with the reported resolutions. The scatter plot indicates high correlation (R^2^=0.91) between the 77 resolution estimates from both approaches (**Figure 2d)**. As expected in the absence of any masking for both approaches, the correlation of the 1% FDR-FSC with the 0.143 FSC threshold deteriorates due to the presence of solvent noise (**Figure 2e).** The resolution differences to the reported EMDB resolutions using user-defined masks are minimal in the case of 1 % FDR-FSC whereas the differences with the unmasked 0.143 values possess a median deviation of 0.5 Å (**Figure 2f**, **Supplementary Table 1)**. Although for lower-resolution structures, deposited half-maps are rare and resolutions are more difficult to validate based on the visible features, results of our 1% FDR-FSC procedure also show good agreement with the reported resolutions in such cases (**Table 1**). In conclusion, the determined resolution estimates using 1 % FDR-FSC are in close agreement with previously reported resolution values from the EMDB and can be computed in a fully automated fashion without any user interference.

**Table 1.**
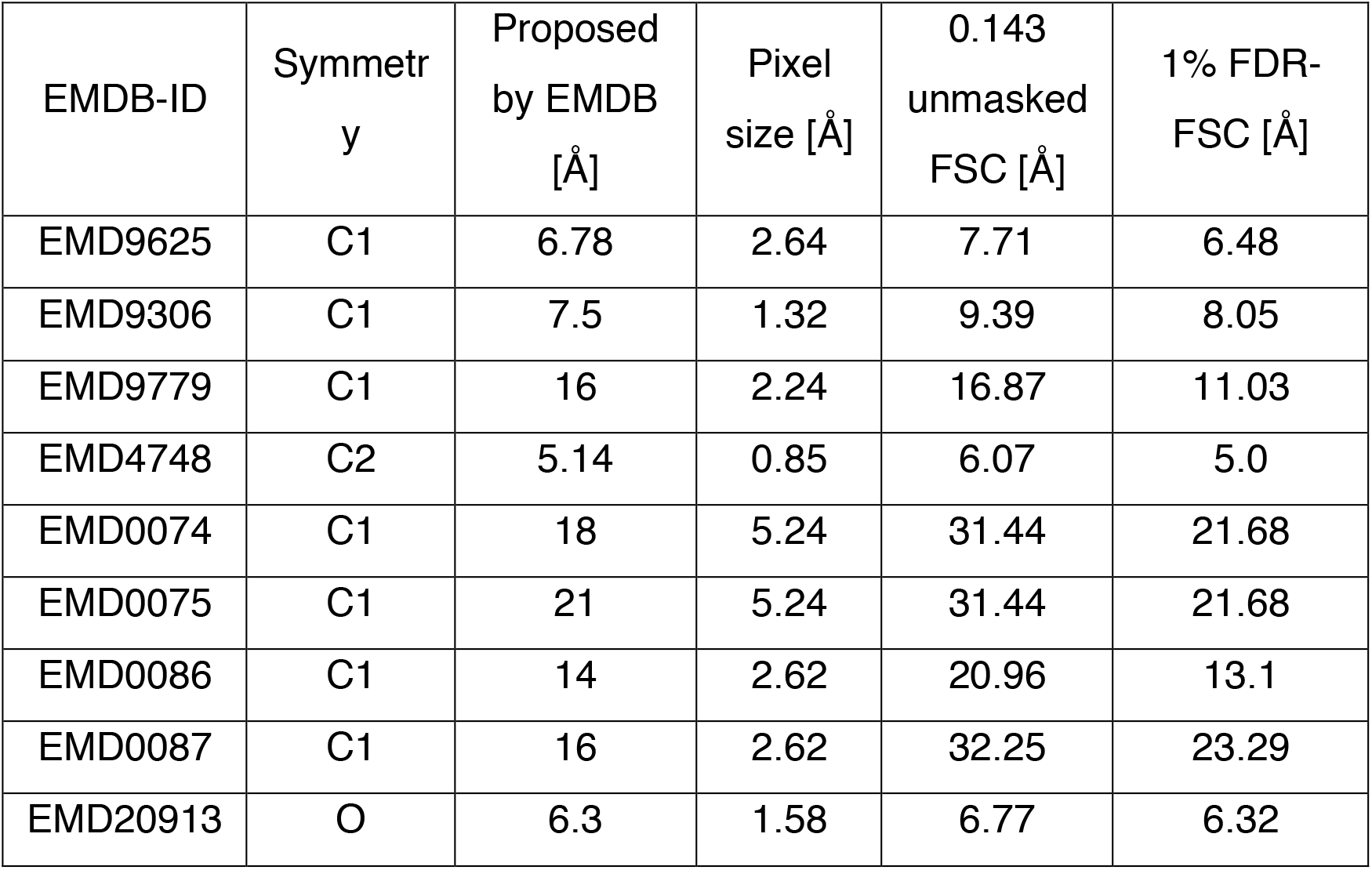
FDR-FSC resolution estimates for low-resolution maps.

| EMDB-ID | Symmetry | Proposed by EMDB [Å] | Pixel size [Å] | 0.143 unmasked FSC [Å] | 1% FDR-FSC [Å] |
| --- | --- | --- | --- | --- | --- |
| EMD9625 | C1 | 6.78 | 2.64 | 7.71 | 6.48 |
| EMD9306 | C1 | 7.5 | 1.32 | 9.39 | 8.05 |
| EMD9779 | C1 | 16 | 2.24 | 16.87 | 11.03 |
| EMD4748 | C2 | 5.14 | 0.85 | 6.07 | 5.0 |
| EMD0074 | C1 | 18 | 5.24 | 31.44 | 21.68 |
| EMD0075 | C1 | 21 | 5.24 | 31.44 | 21.68 |
| EMD0086 | C1 | 14 | 2.62 | 20.96 | 13.1 |
| EMD0087 | C1 | 16 | 2.62 | 32.25 | 23.29 |
| EMD20913 | O | 6.3 | 1.58 | 6.77 | 6.32 |

### Application of FDR-FSC to local resolution estimation

Cryo-EM maps usually exhibit local variations of resolutions and estimating local resolutions has become critical in order to evaluate the structures (Cardone *et al.*, 2013; Kucukelbir *et al.*, 2014; Vilas *et al.*, 2018). We reasoned that the adaptive FDR-FSC thresholding approach could provide benefits over fixed threshold approaches. Therefore, we extended our approach by computing local FSC curves and tested it on cases with large resolution variations. For initial testing, we simulated a composite map from *β*-galactosidase (PDB ID 5a1a(Bartesaghi *et al.*, 2015)) of filtered subunits at resolutions of 2.0, 3.0, 4.0 and 5.0 Å and included Gaussian white noise. Next, we computed the local FSC’s from overlapping cubes including a Hann window of half the cube dimension. Using local 1% FDR-FSC thresholding, we determined local resolutions measures at different noise levels. In comparison, the commonly used 0.5 local FSC threshold shows a stronger deviation at higher noise levels and tends to underestimate resolution in comparison with the 1% FDR-FSC threshold at low noise levels (**Supplementary Figure 5d, e**). Application to an experimental 3.4 Å *γ*-secretase map (Bai *et al.*, 2015)(EMD3061) shows that the 1% FDR-FSC criterion assigns higher resolutions to the best-resolved core of the membrane protein when compared with the 0.5 FSC cutoff, while at the same time, lower resolutions between 10 and 15 Å are found at peripheral glycosylations and the detergent belt (**Figure 3a** **top left**). Although simulated and experimental maps indicated the utility of the approach, one important aspect is the limited number of Fourier coefficients included in the FSC computation as determined by the window size. In order to test the window size effect on the resolution, we computed the local resolutions by the 1% FSCR-FDR as well as the 0.5 local threshold FSC method of the experimental 3.4 Å *γ*-secretase map (Bai *et al.*, 2015)(EMD3061) using increasing window sizes from 10 to 50 pixels (**Supplementary Figure 6a**). For very small windows (10 and 15) in the center, we observe an underestimation of resolution presumably due to undersampling, whereas for windows larger than 20 pixels we observe constant resolution values for two exemplary windows (**Supplementary Figure 6b**). In the transmembrane region of the protein, the resolution is overestimated at 3.8 – 4.0 Å using the local 0.5 method in comparison with the 3.4 Å resolution estimated by the 1 % FDR-FSC method. The latter values are supported visually by the features of the cryo-EM density map. In contrast, peripheral regions next to the protein show high sensitivity to which structural regions are included in the measurement in line with previous observations on FSC-based local resolution measurements (Cardone *et al.*, 2013). Following Cardone and colleagues, we confirm that using window sizes of seven times the overall resolution provides a good parameter for robust local 1% FDR-FSC resolution determination.

**Figure 3.**
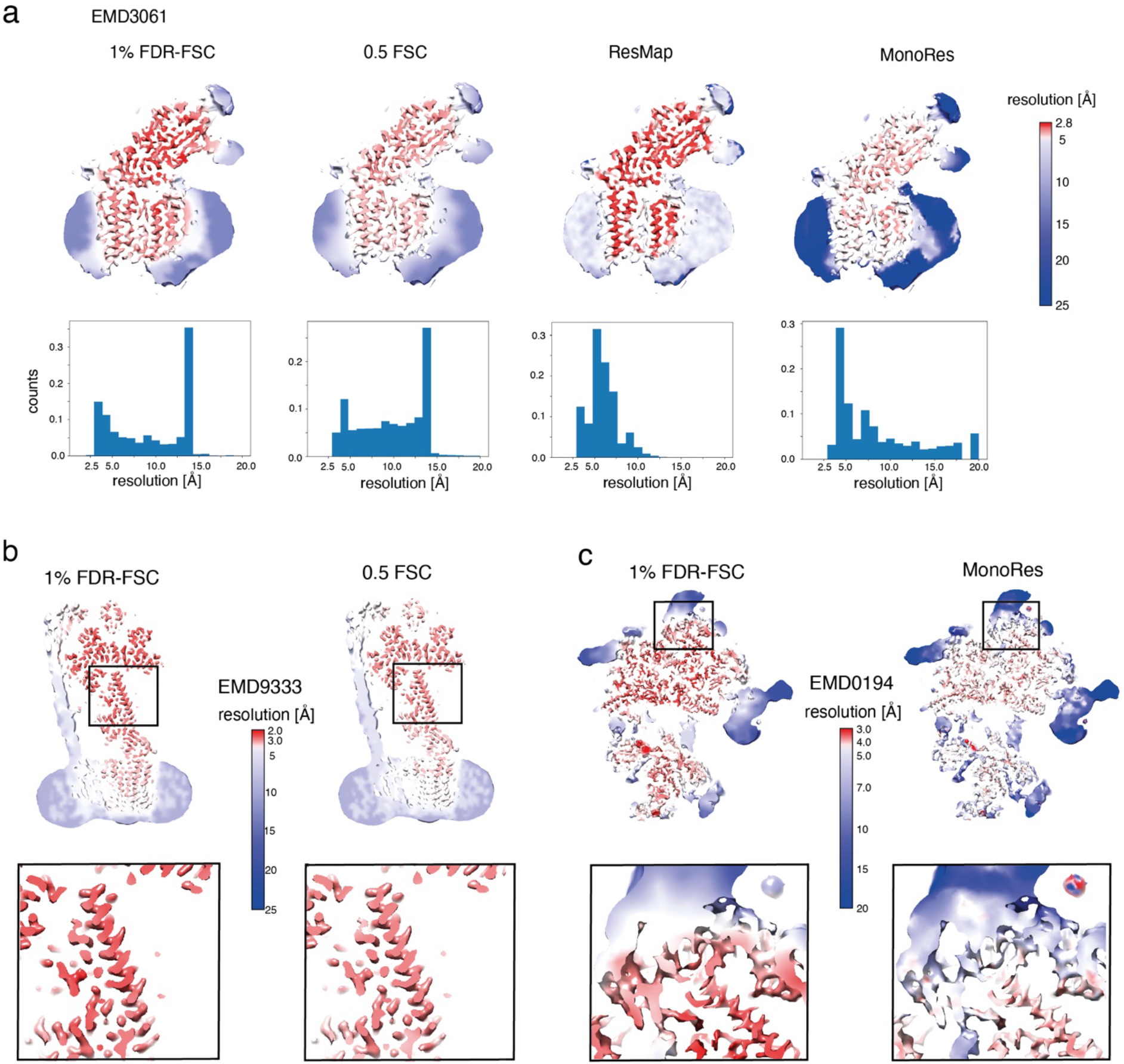
Application of FDR-FSC to local resolution estimation. (a) Comparison of local resolutions estimated for the *γ*-secretase map EMD3061 for 1% FDR-FSC, 0.5 FSC, ResMap and MonoRes grouped with corresponding resolution histograms below. (b) Surface mapped local resolutions determined using 1% FDR-FSC (left) of a 3.9 Å map of a bacterial ATP synthase (EMD9333) compared with 0.5 FSC mapping (right). (c) Surface mapped local resolutions determined using 1% FDR-FSC of a 3.8 Å map of a eukaryotic ribosome (EMD0194) compared with MonoRes mapping.

Further comparisons with available software packages computing local resolution show that using standard settings applied to the *γ*-secretase map, ResMap (Kucukelbir *et al.*, 2014) assigns more optimistic resolutions, whereas MonoRes (Vilas *et al.*, 2018) gives estimates of more conservative resolution (**Figure 3a**). The local resolution histogram of ResMap lacks a low-resolution peak from the detergent whereas MonoRes assigns too conservative estimates around 4-5 Å to the high-resolution parts, similar to 0.5 FSC (**Figure 3a** **bottom**). The local resolution histogram of the 1% FDR-FSC method covers both aspects of high and low-resolution features well. Additional analyses of a 3.9 Å ATP synthase (Guo *et al.*, 2019)(EMD9333) and a eukaryotic ribosome (Juszkiewicz *et al.*, 2018) (EMD0194) map confirms this notion that the 1% FDR-FSC criterion assigns high resolutions to the core map parts while avoiding overestimation of resolution in the low-resolution map parts (**Figure 3b** **and** **3c**). In these cases, the 1% FDR-FSC estimated resolution measures can be validated by visual features appearance such as the low-resolution L1-stalk domain >7 Å and the clearly visible side-chain density of the *α*-helical segment at 3-4 Å. Despite the sensitivity to window sizes inherent to any local FSC determination, the here presented 1% local FDR-FSC approach can robustly assign local resolution in cases of highly variable local noise levels.

### Application of FDR-FSC to directional resolution estimation

In addition to local resolutions, directional resolutions have been recently evaluated to assess the effect of preferred particle orientations (Zi Tan *et al.*, 2017). In practice, the FSC curve from a conical 3D Fourier transform is calculated including voxels of a specified angle. In analogy to local resolution windows, due to the limited number of Fourier coefficients inside a conical volume, directional FSCs suffer from poor statistics and therefore the FDR-FSC approach could provide similar benefits as demonstrated in the case of local FSCs. We tested the approach in more detail using three different cases: a map of the soluble portion of the small influenza hemagglutinin (HA) trimer with highly preferred orientations (EMD8731, **Figure 4a**) (Zi Tan *et al.*, 2017), a highly symmetric apo-ferritin map (EMD0144, **Figure 4b**) (Zivanov *et al.*, 2018) and an asymmetric map of *γ*-secretase (EMD3061, **Figure 4c**) (Bai *et al.*, 2015). Inspection of the directional resolution plots reveals that the 0.143 FSC-thresholds tend to give more optimistic resolution estimates compared with the 1% FDR-FSC approach. This effect is more pronounced at lower resolutions. As a result, the 1% FDR-FSC criterion displays stronger resolution differences with lower resolutions up to 8 Å in the vertical direction and higher resolutions up to 4.2 Å in the horizontal direction (**Figure 4a** **and** **4c**). Due to highly preferred orientations, such a result can be expected in this sample (Zi Tan *et al.*, 2017). In contrast, the directional values derived by the 0.143 FSC threshold are not able to resolve the differences in directional resolution apparent in the HA map. For the 1.6 Å resolution map of apo-ferritin, clear directional resolution differences are not displayed due to high symmetry and homogenous particle orientations (**Figure 4b**). Similarly, the asymmetric map of *γ*-secretase only exhibits minor directional resolution differences (**Figure 4c**). We attribute the larger detected resolution differences by the FDR-FSC approach to the more faithful resolution determination in cases of smaller number of Fourier coefficients in analogy to the benefits observed for local resolution. In conclusion, FDR-FSC is suitable for local resolution determination as well as for directional resolution measurements as it estimates resolution at different noise levels more robustly in comparison with fixed threshold approaches.

**Figure 4.**
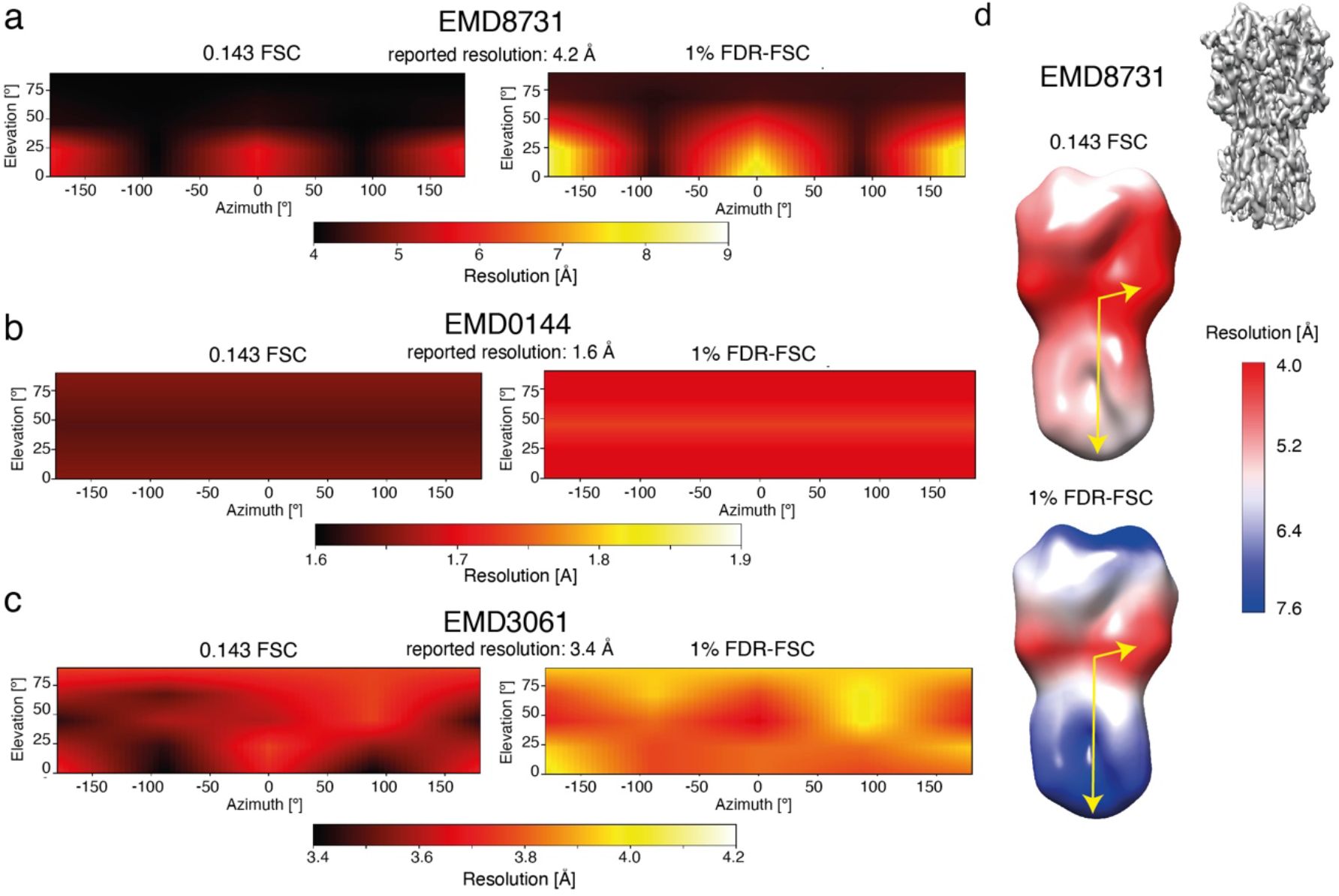
Application of FDR-FSC to directional resolution estimation. Comparison of directional resolution plots of EMDB entries (a) EMD8731, (b) EMD0144 and (c) EMD3061 for 0.143 FSC (left) and 1% FDR-FSC thresholds (right). Resolutions are shown in colors for the respective directions corresponding to angles of azimuth and elevation. (d) Directional resolutions mapped on the low-pass filtered surface of EMD8731. The resolution at each voxel specifies the resolution in the direction given by the vector from the center to the respective voxel (c.f. yellow arrows). Stronger directional resolution differences are reported using the 1% FDR-FSC measurement.

## 3. Discussion

Resolution estimation is one of the essential tasks to assess the experimental quality and confidence for the interpretation of cryo-EM maps. Therefore, an automated procedure with least user interference delivering robust results is desirable. Here, we propose a new approach of thresholding of FSC curves by non-parametric permutation sampling followed by FDR control and show that this approach is suitable for reproducible resolution determination. The approach does not have any free parameters. The only additional information that needs to be supplied is the volume symmetry applied. This way, the FDR-FSC approach enables resolution estimation without masking and thus eliminates the requirement of the mask including the optional deconvolution process (Chen *et al.*, 2013). Although mask-free approaches have been proposed (Sindelar & Grigorieff, 2012), they require estimates of the expected molecular volume, which corresponds to information that may not be routinely available from the cryo-EM experiment in particular when heterogeneity is involved. Further application to local resolution estimation showed that judged by visual map features the adaptive FDR-FSC method captures locally high-resolution features in the core of protein structures as well as lower resolution in the periphery of a protein complex. Complementary to local resolutions, directional FSCs can also be analyzed by FDR-FSC and resulted in well-estimated resolutions in cases of anisotropic reconstructions. The algorithm is implemented in a Python program that takes a few seconds for small maps (<200 pixels volume size) up to a few minutes for big maps (>400 pixels volume size).

One potential conceptual advantage of the proposed FDR-FSC approach is that the method only assesses whether any significant correlations beyond random noise can be detected in a given resolution shell. This is different to the commonly used approach of determining the absolute spectral information content of the structure of interest. In this way, the FDR-FSC approach tolerates even high noise levels (see **Supplementary Figure 5b/c**) and therefore does not require any solvent masking. In addition, inference of statistically significant signal in the resolution shells only requires the distribution of random noise correlations determined by permutation and avoids the consideration of complicated correlations between signal and noise (Heel & Schatz, 2017). Once spectral noise distributions have been estimated, *p*-values of the measured FSC value for each resolution shell can be derived. Finally, FDR control of these *p*-values results in the final thresholding step. FDR control is a statistically well-established routine for identifying thresholds while taking into account arbitrary dependencies and varied number of samples (Benjamini & Yekutieli, 2001). Both of these issues occur in FSCs computed from cryo-EM maps. First, correlations and dependencies between neighboring resolution shells in cryo-EM are known to exist due to uncertainties in alignment and are propagated to the reconstruction volume. Second, limited number of Fourier coefficients are present in smaller windows in cases of FSC-based local and directional resolution estimation. The FDR control procedure automatically adjusts the effective *p*-value threshold to the number of available resolution shells. For such applications, however, we also showed that the window size remains a free parameter that affects the computation of local resolutions regardless of which thresholding procedure is being used (see **Supplementary Figure 6**). This is a general property when local FSC calculations are computed and is independent from the respective threshold criterion as it is already elaborated in detail in (Cardone *et al.*, 2013). Irrespective of the window size, the here presented FDR-FSC method has advantages over 0.5 FSC, as it is less sensitive to differences in noise levels in the center and periphery of the protein that can affect the resolution determination.

Given its robustness and minimal input requirement, it is conceivable that the proposed FDR-FDR approach can be used to re-evaluate many deposited EMDB structures when raw half-maps were available. In our evaluation, we found very good agreement between the reported resolution values, mostly based on the 0.143 FSC criterion, and the FDR-FSC resolution values. This correlation shows that the overwhelming majority of depositors submits their resolution value with care in correspondence to the presented visual features such as beta-strand separation and side-chain densities. For future EMDB depositions, the FDR-FSC method could be used as an additional resolution assessment in the context of other standard validation tools. Due to the increased statistical robustness, we see particular use for extending resolution determination into more challenging applications such as local and directional resolution where resolution reporting is often critical for interpretation but technically less standardized. In the current manuscript, we presented the FDR-FSC method with a focus on cryo-EM map evaluation. It has to be noted that the proposed framework is applicable to 2D Fourier ring correlations as used in super-resolution microscopy (Nieuwenhuizen *et al.*, 2013). Due to the demonstrated ease of use and robustness, we anticipate a common applicability of the FDR-FSC method to image analysis and processing tools in and beyond the cryo-EM field.

## 4. Methods

### 4.1 Permutation testing for Fourier shell correlation coefficients

Let 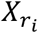 and 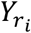 complex random variables and we denote the corresponding Fourier coefficients at the specific location *r*_*i*_, *i* = 1, … , *N*, in resolution shell *r* of half-map 1 and 2, where *N* ∈ ℕ is the number of Fourier coefficients in the respective shell. Due to the Friedel symmetry inherent to Fourier transforms, we restrict ourselves to one half of all locations *r*_*i*_. The pair is simply represented as the complex conjugate. We denote the Friedel symmetry corrected sample size with *n*. In the case of two independent reconstructions, dependence between 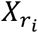and 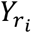 is introduced through an effect of the signal at position *r*_*i*_, which we denote with 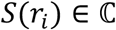 and which is the same in both volumes. Thus, Fourier coefficients are usually modelled as

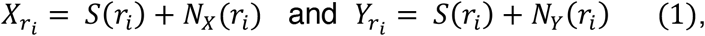

where *N*_*X*_(*r*_*i*_) and *N*_*Y*_(*r*_*i*_) are complex valued noise variables (Van Heel & Schatz, 2005). No assumptions about the distribution of the noise in resolution shell *r* are made. Furthermore, we denote by 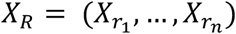 and similarly 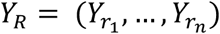 the complete set of resolution shells.

Initially, we have to clarify whether the Fourier coefficients in the respective resolution shell are dependent with respect to the location *r*_*i*_, i.e. that groupings 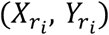 show statistical dependencies. Expressed in statistical terms: the null hypothesis *H*_0_ is that 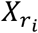 and 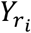 independent of each other, and the alternative hypothesis *H*_1_ is that there are dependencies. In order to test such a hypothesis, correlation coefficients can be used (Lehmann & Romano, 2005). As a test statistic we use the Fourier shell correlation. Using the Friedel symmetry inherent to Fourier transforms, we write the *FSC* as

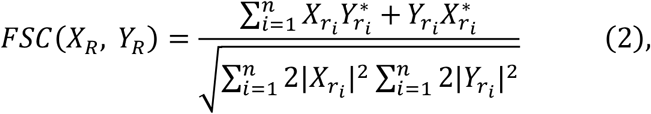

and it becomes clear that *FSC* (*X*_*R*_, *Y*_*R*_) is a real valued random variable. Statistically, *FSC* (*X*_*R*_, *Y*_*R*_) is the estimator of the true value of the Fourier shell correlation, which we denote from now on with FSC. It is important to emphasize here that we are testing for dependencies by using a correlation coefficient; i.e. the null hypothesis is not FSC = 0.

In order to test *H*_0_, we calculate a *p*-value as follows. Using *fsc* we denote the estimated value for the FSC in resolution shell *r*. Moreover, using *FSC* we denote the random variable describing the FSC as above. The *p*-value *p* of this observation is then given as the probability that *FSC* is bigger than *fsc* under the null hypothesis, i.e.

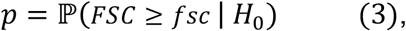

where 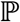 is the true probability distribution under the null hypothesis.

A permutation test is performed as follows. Under *H*_0_, the Fourier coefficients 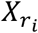 and 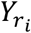 are independent for *i* = 1, … , *n*. Newly paired samples of Fourier coefficients can be generated by permutation of *Y*_*R*_. Thus, the sampled null distribution is the distribution of *FSC* from independent half-maps. For a detailed discussion regarding permutation of correlation coefficients we refer to (DiCiccio & Romano, 2017). From each generated sample, we then calculate the Fourier shell correlation, which results in a sample of *FSC* under the null hypothesis. Denoting with *S*_*n*_ the set of all permutations *π* of {1, … , *n*}, we can calculate the *p*-value *p* by

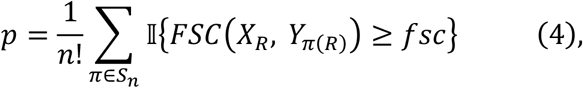

where 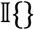 denotes the indicator function. As the number of possible permutations grows rapidly with the sample size, technically often just a random subset *H* ⊂ *S*_n_of all *n*! permutations is used. Thus, *p* is replaced by its Monte-Carlo estimator 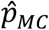, given as

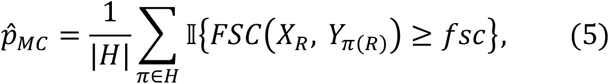

where |*H*| is the cardinality of the set *H*, i.e. the number of permutations selected for the Monte-Carlo estimate. It is important to note that the null distribution of the *FSC* estimated by the permutation approach is not necessarily the distribution of the *FSC* in the absence of any signal. As we are permuting in the presence of possible signal, which adds additional variation, the permutation distribution of the *FSC* will have larger tail probabilities than the *FSC* distribution without any signal. Consequently, this could give rise to more conservative resolution estimates. However, simulations in the presence of signal at high and low signal-to-noise ratios showed that this effect seems to have less practical relevance (**Supplementary Figure 7**).Moreover, the amount of signal in the most critical resolution shells close to the true resolution of the structure is low and will thus have limited influence on the actual distribution. As a compromise between computational efficiency and statistical accuracy, we restrict the number of permutations to a maximum of 1,000. Permutations are only performed for resolution shells with an effective sample size >10, which allows for more than 1,000 permutations. In cases of insufficient sampling below 10, the program will generate a warning message.

### 4.2 Multiple testing correction

Testing the complete set of resolution shells of a volume results in a multiple testing problem. In order to correct for the multiple testing problem, estimated *p*-values are subsequently adjusted for the false discovery rate (FDR) of detected resolution shells, i.e. the maximum amount of false positive resolution shells. The FDR is given as

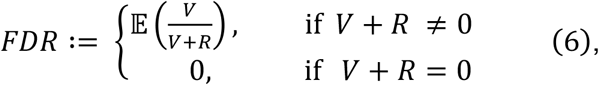

with *V* ∈ ℕ_0_ the number of false rejections, *R* ∈ ℕ_0_ the number of true rejections and 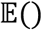 denotes the expectation value. For a more detailed description of the exact adjustment in the context of cryo-EM we refer to (Beckers *et al.*, 2019). The false discovery rate (FDR) is controlled with the Benjamini-Yekutieli procedure (Benjamini & Yekutieli, 2001), which controls the FDR under arbitrary dependencies.

### 4.3 Effective sample size corrections

Correction of the sample size has been found to be an important factor in presence of symmetry and masking (Van Heel & Schatz, 2005) (Results section), as this leads to additional dependencies between Fourier coefficients and thereby reduces the true sample size to an effective sample size *n*_*eff*_. The effect of imposed symmetry during reconstruction can be conservatively corrected by

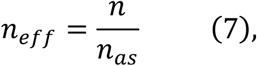

where *n* is the number of Fourier coefficients in the respective resolution shell and *n*_*as*_ is the number asymmetric volume units at the given symmetry (**Supplementary Figure 1b**). Effective sample sizes are incorporated in the permutation framework by sub-sampling of Fourier coefficients. Effects of masking on the effective sample size are more complicated and depend on the specific shape and volume of the mask. Here, we estimate the effective sample size *n*_*eff*_ ≈ *α* * *n* by finding the factor *α* ∈ (0,1] through minimization of the mean Kolmogorov-Smirnov distance 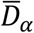 over the resolution shells. The two-sample Kolmogorov-Smirnov statistics (Massey, 1951) *D*_*α*_ is a measure of similarity of two empirical cumulative distribution functions (ECDF) and is defined as

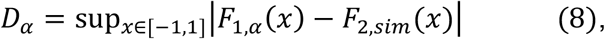

where in our setting *F*_1,*α*_(*x*) denotes the ECDF of the permutation approach. This approach is estimated using an effective sample size *N*_*eff*_ ≈ *α* ∗ *n*, and *F*_2,*sim*_(*x*) an estimate of the true cumulative distribution function in presence of the respective mask. *F*_2,*sim*_(*x*) can be obtained by repetitive simulation of two masked noise maps and subsequent FSC calculation. 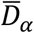 is then calculated as the mean of the Kolmogorov-Smirnov statistics *D*_*α,r*_ of the resolution shells *r*, i.e.

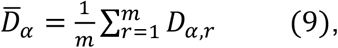

where *m* is the number of resolution shells. An estimate 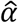 for *α* is then given by

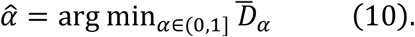

In the presented algorithm, we apply a soft circular mask for global resolution estimation by default, as the resulting effect can be corrected by an effective sample size of *n*_*eff*_ ≈ 0.7*n*, i.e. 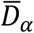 was minimized for *α* ≈ 0.7 (**Supplementary Figure 2a**), which allows accurate calculation of tail probabilities for various box sizes (**Supplementary Figure 3**). Application of windowing functions, as used for local resolution estimation (see below), also leads to reduced effective sample sizes. In a similar way as for the soft circular mask, we found that a Hann-window leads to an effective sample size *n*_*eff*_ ≈ 0.23*n* (**Supplementary Figure 2b**, **Supplementary Figure 4**).

### 4.4 Local Resolution estimation

Local resolutions are estimated by a moving window across both half maps and subsequent calculation of local FSC thresholds (Cardone *et al.*, 2013). In order to account for high-resolution artifacts introduced through spectral leakage, a Hann-function is used as a windowing function. The masking effects, which are introduced from the Hann window, are corrected as described above with a correction factor *α* ≈ 0.23. Moreover, effects of symmetry do not have any influence on the effective sample sizes when the size of the sliding windows is smaller than the size of the asymmetric volume unit in the map. In order to speed up the calculations, a step-size option is implemented allowing movement of the sliding window of more than a single voxel. Moreover, in order to avoid repeating permutations on the same map, the permutations are only done on 10 random locations of the sliding window. The resulting samples of the null distribution are merged and subsequently used for *p*-value calculation at all locations. We found that window sizes around seven times the estimated global resolution provide a reasonable compromise between locality and resolution sampling in Fourier space for most cases. Presented experiments were performed using a window size of 25 pixels.

### 4.5 Directional resolution estimation

The implementation of the directional resolution estimation follows Lyumkis and colleagues (Zi Tan *et al.*, 2017). For each direction, the FSC is calculated by taking those voxels from each half-map that are included by a specified angle at the respective direction. This results in rotating an inverse cone over one half of the 3D Fourier transforms, thereby accounting for the Friedel symmetry, and calculating the FSC only from samples inside the inverse cone. In analogy to (Zi Tan *et al.*, 2017), an angle of 20° was used for the presented experiments providing a good compromise between the number of Fourier coefficients per shell and the preservation of local directionality. Sample size corrections need to be applied as in the case of local resolution. Similar to global resolution estimation, this results in a correction factor *α* ≈ 0.7 due to soft spherical masking. Moreover, in analogy to local resolutions, locality of the directional resolution does not require the correction of symmetry effects. As implemented in the case of local resolution estimation, in order to accelerate the algorithm, the resolutions are only calculated for a limited set of directions and results are interpolated in order to avoid repetitive FSC calculations of overlapping cones.

### 4.6 Software

The procedures are implemented together with the previously published Confidence Map tools in a GUI based software named SPoC – Statistical Processing of cryo-EM maps (**Supplementary Figure 8**). Code is written in Python3 based on NumPy (Van Der Walt *et al.*, 2011), matplotlib (Hunter, 2007), SciPy (Oliphant, 2007), mrcfile (Burnley *et al.*, 2017) and parallelized with multiprocessing. The software is available at https://github.com/MaximilianBeckers/SPOC. Figures were prepared with UCSF Chimera (Pettersen *et al.*, 2004).

## Author contributions

M.B. and C.S. designed the research. M.B. developed and implemented the algorithm and conducted the experiments. M.B. and C.S. wrote the manuscript.

## Acknowledgements

We are grateful to Thomas Hoffmann and Jurij Pecar (IT Services) for maintenance of the high-performance computing at EMBL.

---

**Supplementary Figure 1.**
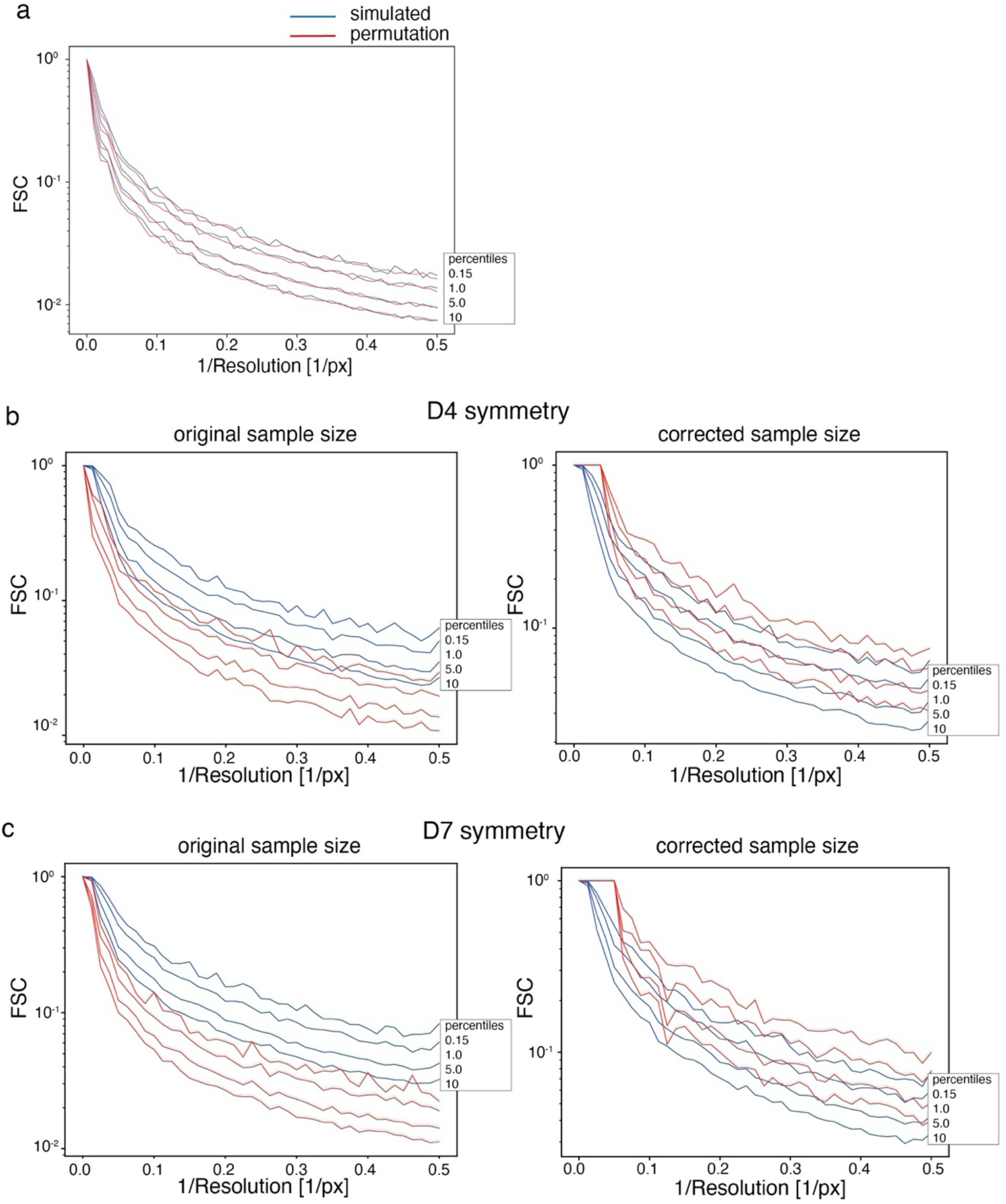
Comparison of permuted and simulated half-map FSC distribution and the effect of symmetry on sample size and FSC distributions. (a) Right-sided 10, 5.0, 1.0 and 0.15 percentiles obtained from FSC permutations within a resolution shell (red), and from 5,000 noise-simulated FSCs (blue). The y-axis corresponding to the FSC values is formatted on a logarithmic scale. (b) Comparison of the respective percentiles between the permutation approach and simulation of the true FSC distribution for the case of imposing (b) D4 and (c) D7 symmetries. Plots on the left side show that the presence of symmetry reduces the effective sample size, which leads to underestimated probabilities at the tail, and can be conservatively corrected (right). The y-axis corresponding to the FSC values is formatted on a logarithmic scale.

**Supplementary Figure 2.**
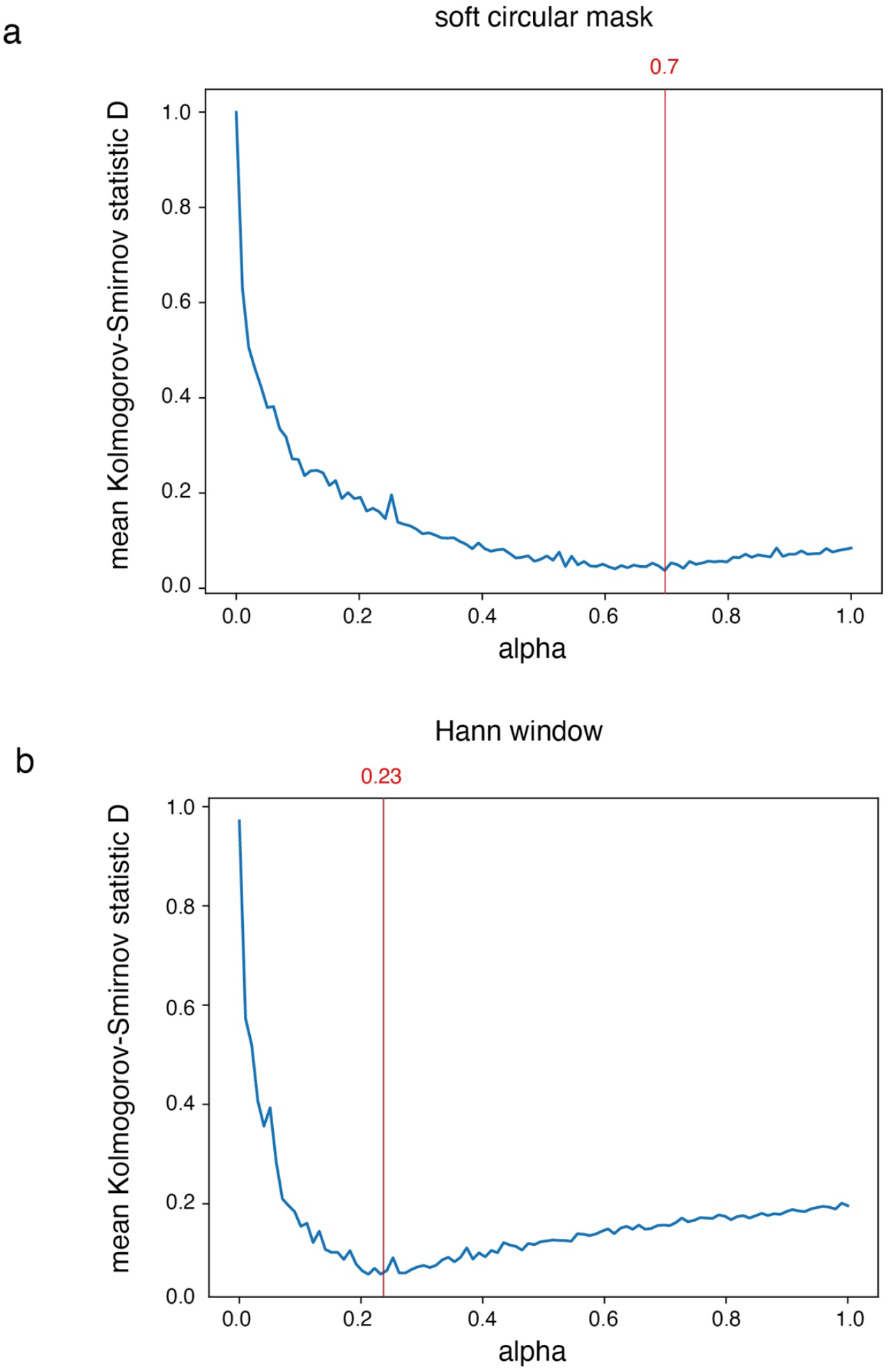
Estimation of sample size correction factor. Sample size correction factor *α* plotted against the mean Kolmogorov-Smirnov statistic distance of the distributions of the resolution shells. (a) For a soft circular mask the minimal distance of *α* is ≈ 0.7. (b) For a Hann window the minimum distance for *α* corresponds to ≈ 0.23.

**Supplementary Figure 3.**
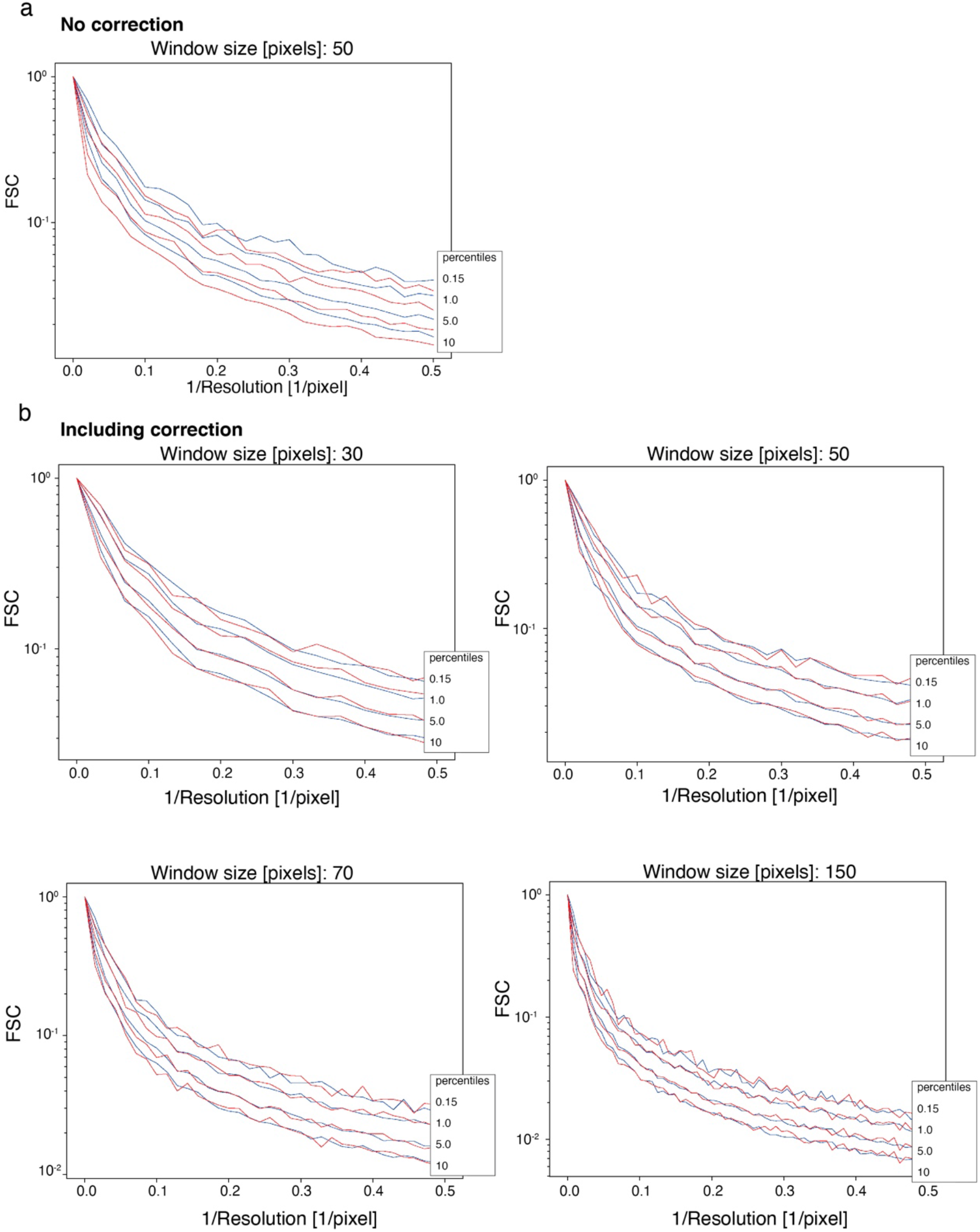
Effective sample size after circular masking. Comparison of right-sided 10, 5.0, 1.0 and 0.15 percentiles, as estimated by permutation (red lines), with the true FSC distribution (blue lines), as obtained from 5,000 simulations of the respective noise half-maps. Here, the influence of a soft circular mask is tested. (a) No correction for the masking effect, distributions at the tail are underestimated. (b) This can be well corrected with an effective sample of ~0.7 for different window sizes. The y-axis corresponding to the FSC values is shown in a logarithmic scale.

**Supplementary Figure 4.**
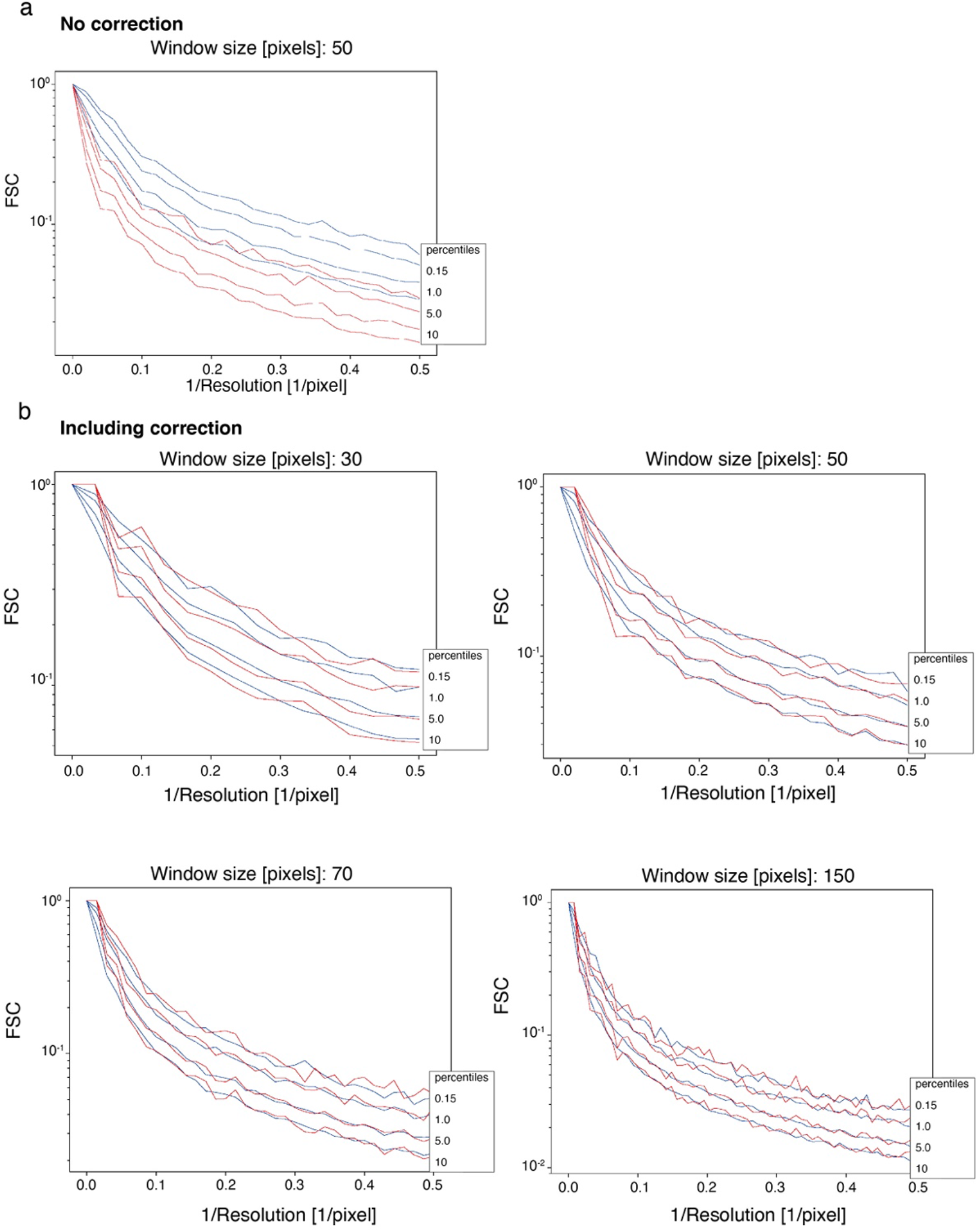
Effective sample size after application of a Hann window. Comparison of right-sided 10, 5.0, 1.0 and 0.15 percentiles, as estimated by permutation (red lines), with the true FSC distribution (blue lines), as obtained from 5,000 simulations of the respective noise half-maps. (a) No correction for the masking effect, the distributions at the tail are underestimated. (b) This can be well corrected with an effective sample of ~0.23 for different window sizes. The y-axis corresponding to the FSC values is shown in a logarithmic scale.

**Supplementary Figure 5.**
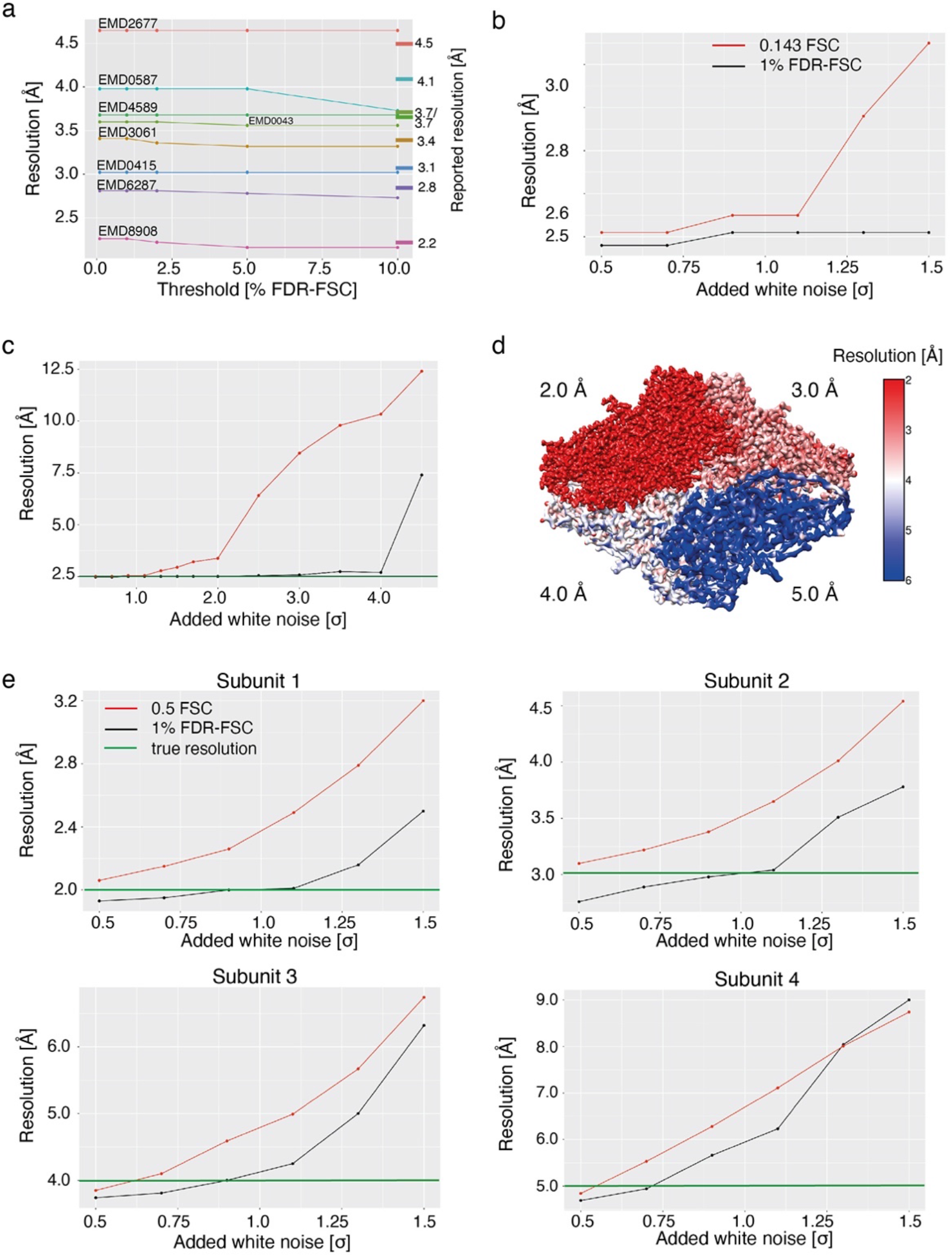
Comparison of FDR-FSC with common fixed threshold FSC resolution determination. (a) Resolutions determined at different FDR thresholds for eight EMDB maps (reported resolution referenced at the right) between 2.2 Å and 4.6 Å resolution show that the FDR threshold is stable towards the specific significance level for reasonable errors levels between 0.1 and 10%. (b and c) Resolution estimates for 0.143 FSC and 1% FDR-FSC thresholds for a simulated map of β-galactosidase (PDB ID 5a1a) at 2.5 Å resolution and with different levels of added background noise up to 4.0 standard deviations. (d) Local resolution estimates of the 1% FDR-FSC threshold for a simulated map of β-galactosidase (PDB ID 5a1a). The 4 subunits were filtered to different resolutions of 2.0, 3.0, 4.0 and 5.0 Å, respectively. (e) Resolution estimates for 0.143 FSC and 1% FDR-FSC thresholds for the map simulated in (d), shown are the mean resolutions for subunit 1 at 2.0 Å, 2 at 3.0 Å, 3 at 4.0 Å and 5 at 5.0 Å from the local resolution measurement. Local resolutions from the 0.5 FSC threshold (red) are compared to 1% FDR-FSC (black). The true resolution of the subunit is referenced in green.

**Supplementary Figure 6.**
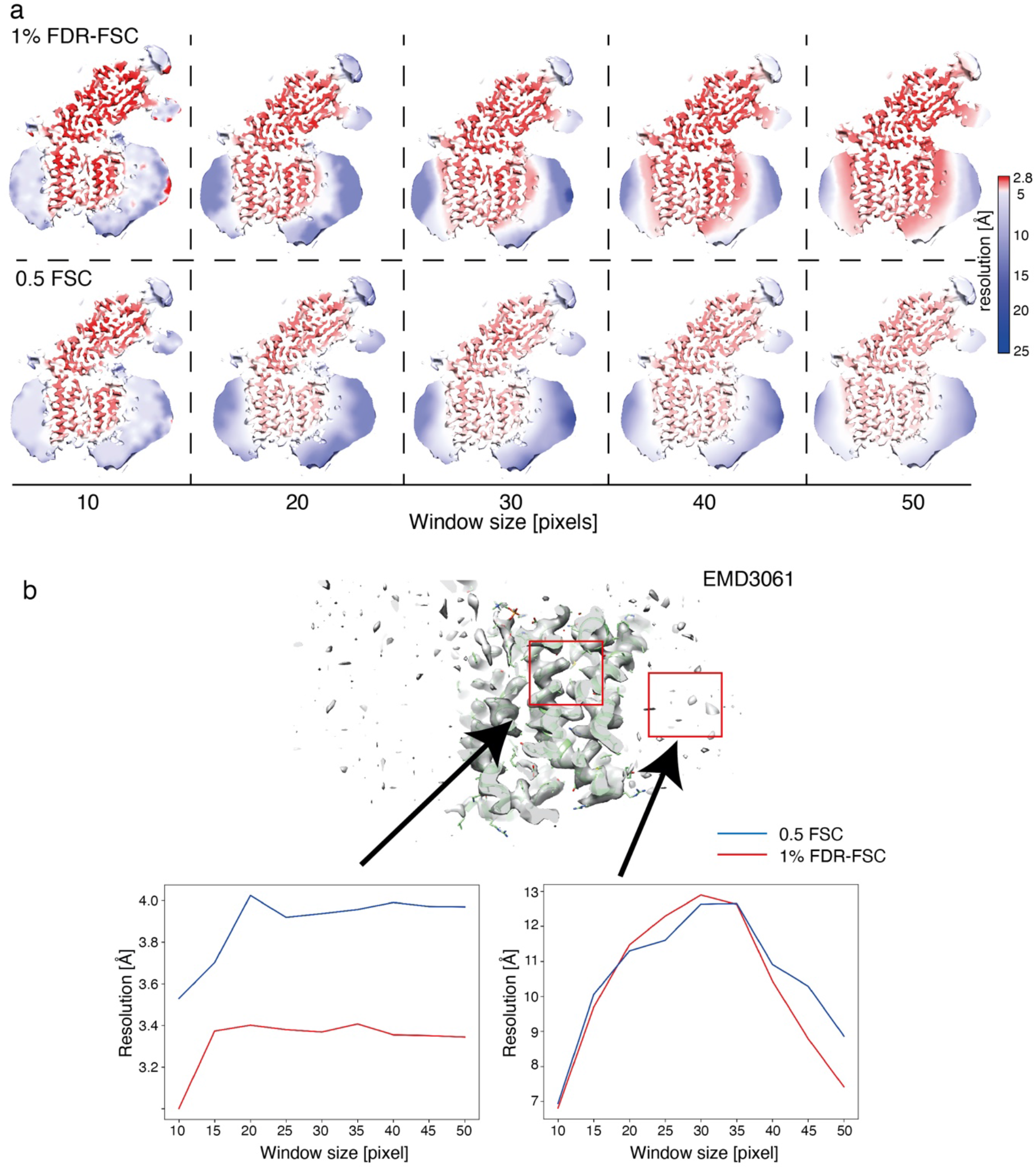
Influence of the window size on local resolutions of the cryo-EM map EMD3061. (a) The local resolutions are mapped on the surface in color code for both 0.5 FSC thresholding and 1% FDR-FSC for window sizes from 10 to 50 pixels. (b) Effect of window size on the local resolutions of a high and a low-resolution region (marked in red in the cryo-EM map).

**Supplementary Figure 7.**
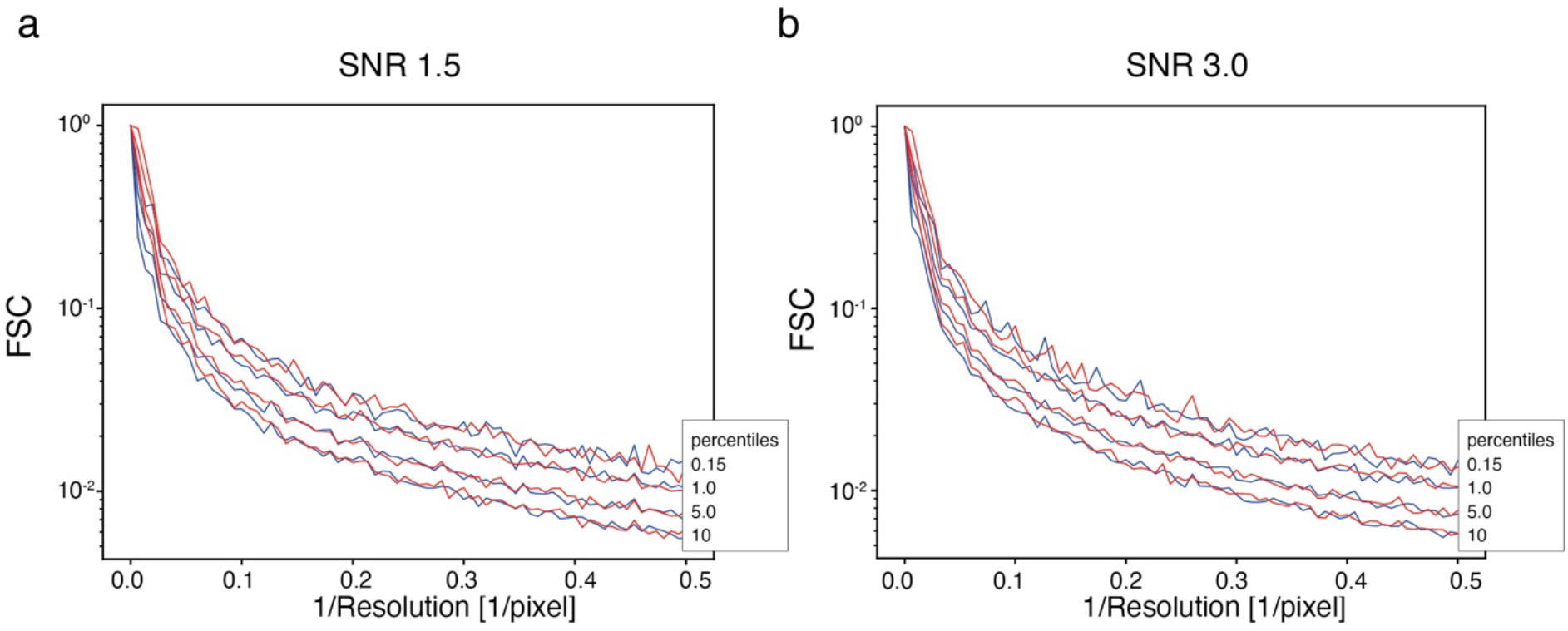
Permutation sampling in the presence of signal at different signal-to-noise ratios. Comparison of right-sided 10, 5.0, 1.0 and 0.15 percentiles, as estimated by permutation (red lines), with the true FSC distribution (blue lines), as obtained from 5,000 simulations of the respective noise half-maps. Here, the influence of present signal on the permutation is tested. Comparison in the presence of signal at a signal-to-noise ratio of (a) 1.5 and (b) 3.0.

**Supplementary Figure 8.**
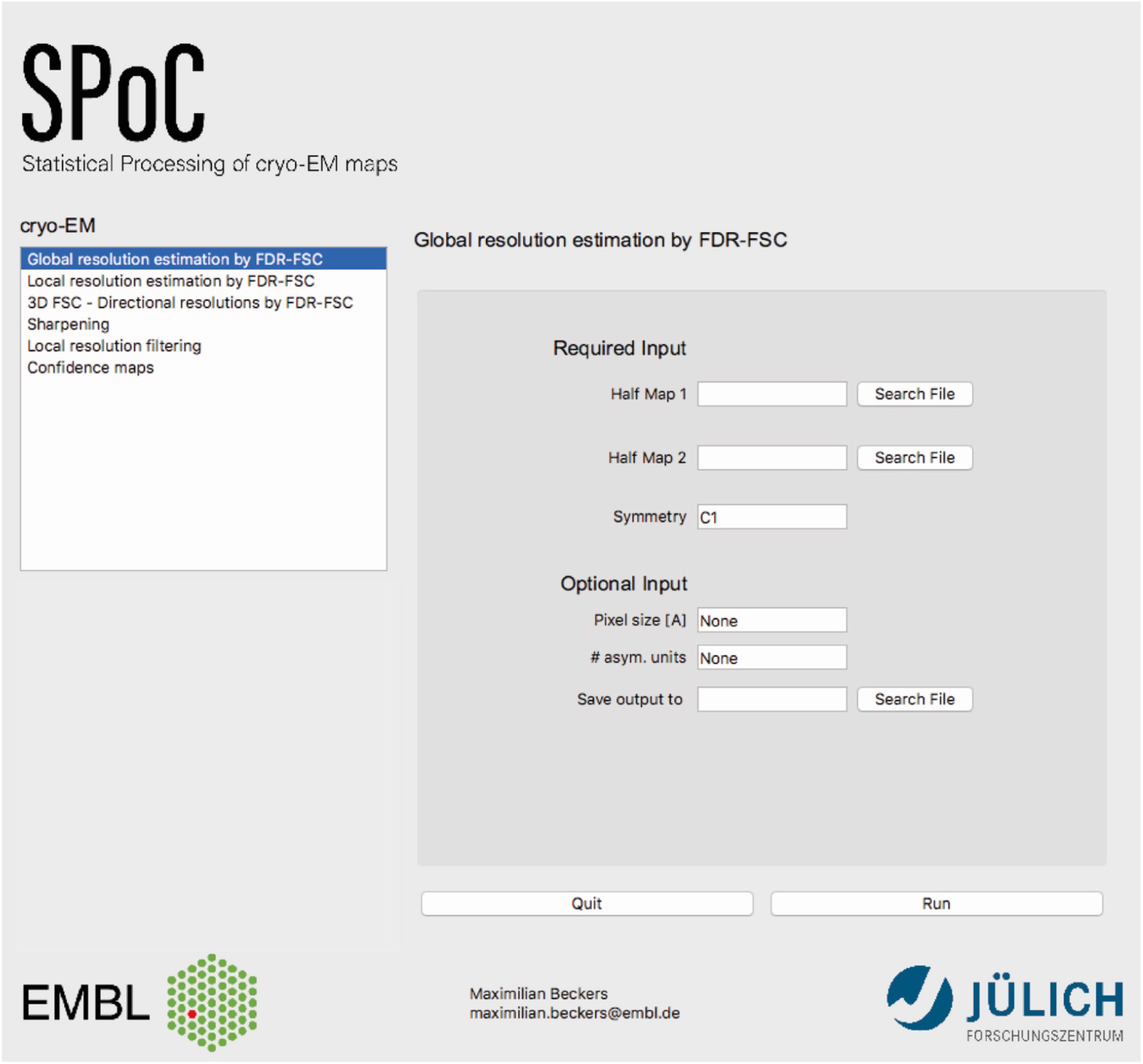
GUI window of SPoC. The presented tools are implemented in an easy-to-use GUI-based software package called SPoC (Statistical processing of cryo-EM maps).

**Supplementary Table 1.** FDR-FSC resolution estimates of 77 maps from the EMDB with reported resolutions better than 5Å.

| EMDB-ID | Symmetry | Proposed by EMDB [Å] | Pixel size [Å] | 0.143 unmasked FSC [Å] | 1% FDR-FSC [Å] |
| --- | --- | --- | --- | --- | --- |
| EMD2677 | C1 | 4.5 | 1.76 | 6.48 | 4.75 |
| EMD3061 | C1 | 3.4 | 1.4 | 4.06 | 3.41 |
| EMD2984 | D2 | 2.2 | 0.637 | 2.74 | 2.51 |
| EMD5778 | C4 | 3.3 | 1.2156 | 3.99 | 3.5 |
| EMD6422 | D7 | 4.1 | 1.07 | 4.51 | 4.28 |
| EMD6287 | D7 | 2.8 | 0.982 | 3.07 | 2.83 |
| EMD5623 | D7 | 3.3 | 1.22 | 3.31 | 3.05 |
| EMD6000 | I | 3.8 | 0.99 | 4.12 | 4.04 |
| EMD2847 | C1 | 2.9 | 0.76 | 3.3 | 2.96 |
| EMD0144 | O | 1.65 | 0.84 | 1.83 | 1.71 |
| EMD0408 | C2 | 3.2 | 0.558 | 3.33 | 2.92 |
| EMD6479 | C1 | 3.5 | 1.31 | 3.57 | 3.23 |
| EMD2764 | C1 | 3.75 | 1.34 | 4.9 | 3.59 |
| EMD0468 | C1 | 3.9 | 1.06 | 6.17 | 4.52 |
| EMD0480 | C1 | 3.54 | 1.06 | 4.31 | 4.13 |
| EMD9384 | C2 | 2.96 | 1.06 | 3.49 | 3.36 |
| EMD0043 | C1 | 3.7 | 1.35 | 3.86 | 3.6 |
| EMD0501 | C1 | 3.5 | 1.00 | 4.17 | 3.47 |
| EMD0500 | C1 | 3.4 | 1.00 | 4.03 | 3.38 |
| EMD0489 | C4 | 4.3 | 1.08 | 5.03 | 4.25 |
| EMD0487 | C4 | 4.0 | 1.08 | 4.25 | 3.89 |
| EMD9333 | C1 | 3.0 | 1.06 | 3.43 | 2.98 |
| EMD0488 | C4 | 3.4 | 1.08 | 3.74 | 3.46 |
| EMD0415 | C1 | 3.1 | 1.067 | 3.84 | 3.02 |
| EMD0416 | C1 | 3.6 | 1.067 | 3.96 | 3.59 |
| EMD0417 | C1 | 3.7 | 1.067 | 4.13 | 3.73 |
| EMD0418 | C1 | 3.8 | 1.067 | 4.52 | 3.96 |
| EMD0419 | C1 | 3.3 | 1.067 | 4.18 | 3.2 |
| EMD0420 | C1 | 3.4 | 1.067 | 4.32 | 3.23 |
| EMD0425 | C1 | 3.5 | 1.067 | 6.31 | 3.42 |
| EMD4587 | C2 | 3.8 | 1.012 | 4.26 | 3.86 |
| EMD4588 | C2 | 3.6 | 1.012 | 3.98 | 3.59 |
| EMD4589 | C2 | 3.7 | 1.012 | 4.12 | 3.68 |
| EMD4592 | C2 | 3.6 | 1.012 | 4.12 | 3.63 |
| EMD4593 | C2 | 3.7 | 1.012 | 4.26 | 3.52 |
| EMD4594 | C2 | 3.6 | 1.012 | 4.19 | 3.74 |
| EMD4611 | C2 | 3.2 | 1.012 | 3.53 | 3.18 |
| EMD4612 | C2 | 3.8 | 1.012 | 3.91 | 3.43 |
| EMD4613 | C2 | 3.5 | 1.012 | 3.75 | 3.36 |
| EMD4614 | C2 | 3.3 | 1.012 | 3.53 | 2.93 |
| EMD9599 | O | 1.62 | 0.52 | 1.77 | 1.68 |
| EMD9012 | I | 1.86 | 0.39 | 1.92 | 1.83 |
| EMD0153 | D2 | 1.89 | 0.637 | 2.29 | 1.91 |
| EMD9203 | D3 | 2.1 | 0.66 | 2.33 | 2.04 |
| EMD7541 | D2 | 2.4 | 0.86 | 3 | 2.77 |
| EMD8908 | D2 | 2.2 | 0.637 | 2.57 | 2.26 |
| EMD4129 | C1 | 3.06 | 1.04 | 3.75 | 2.95 |
| EMD0194 | C1 | 3.8 | 1.52 | 4.33 | 3.45 |
| EMD0407 | C2 | 2.8 | 0.558 | 3.66 | 3.25 |
| EMD0452 | C1 | 3.7 | 1.15 | 4.27 | 3.59 |
| EMD0587 | C1 | 4.1 | 1.15 | 4.83 | 3.98 |
| EMD0588 | C1 | 4.3 | 1.15 | 6.01 | 4.27 |
| EMD0589 | C1 | 3.9 | 1.15 | 4.46 | 3.68 |
| EMD0590 | C2 | 3.13 | 1.09 | 3.63 | 3.19 |
| EMD0591 | C2 | 3.37 | 1.09 | 3.98 | 3.4 |
| EMD0592 | C2 | 3.15 | 1.09 | 3.77 | 3.19 |
| EMD9241 | C1 | 3.8 | 1.34 | 4.47 | 3.6 |
| EMD0182 | I | 3.36 | 1.063 | 3.51 | 3.3 |
| EMD0596 | C1 | 3.08 | 1.150 | 3.69 | 2.75 |
| EMD0183 | I | 3.18 | 1.063 | 3.47 | 3.26 |
| EMD7335 | C1 | 3.5 | 0.84 | 4.17 | 3.51 |
| EMD7334 | C1 | 3.9 | 0.84 | 4.4 | 3.9 |
| EMD7337 | C1 | 4.6 | 2.08 | 5.99 | 4.47 |
| EMD7770 | D2 | 1.9 | 0.637 | 2.13 | 1.84 |
| EMD0264 | C5 | 4.6 | 1.24 | 6.46 | 4.46 |
| EMD9013 | C1 | 3.4 | 0.79 | 4.04 | 3.61 |
| EMD9014 | C1 | 3.3 | 0.79 | 3.49 | 3.11 |
| EMD8731 | C3 | 4.2 | 1.31 | 5.41 | 4.3 |
| EMD0341 | C2 | 3.6 | 1.2 | 4.75 | 3.96 |
| EMD0342 | C2 | 4.2 | 1.2 | 6.17 | 4.28 |
| EMD9696 | C1 | 3.76 | 0.875 | 4.17 | 3.72 |
| EMD9695 | C1 | 3.64 | 0.875 | 4.07 | 3.54 |
| EMD0498 | C6 | 2.7 | 1.07 | 2.85 | 2.45 |
| EMD0499 | C6 | 2.7 | 1.07 | 2.81 | 2.56 |
| EMD20074 | C3 | 4.1 | 1.02 | 4.46 | 4.08 |
| EMD20101 | C1 | 4.36 | 1.09 | 4.81 | 3.93 |
| EMD20100 | C1 | 3.28 | 1.09 | 3.67 | 3.17 |

## References

Bai, X. C., Yan, C., Yang, G., Lu, P., Ma, D., Sun, L., Zhou, R., Scheres, S. H. W. & Shi, Y. (2015). Nature. 525, 212–217.

Banterle, N., Bui, K. H., Lemke, E. A. & Beck, M. (2013). J. Struct. Biol. 183, 363–367.

Bartesaghi, A., Merk, A., Banerjee, S., Matthies, D., Wu, X., Milne, J. L. S. & Subramaniam, S. (2015). Science (80-.). 348, 1147–1151.

Beckers, M., Jakobi, A. J. & Sachse, C. (2019). IUCrJ. 6, 18–33.

Benjamini, Y. & Hochberg, Y. (1995). J. R. Stat. Soc. B. 57, 289–300.

Benjamini, Y. & Yekutieli, D. (2001). Ann. Stat. 29, 1165–1188.

Burnley, T., Palmer, C. M. & Winn, M. (2017). Acta Crystallogr. Sect. D Struct. Biol. 73, 469–477.

Cardone, G., Heymann, J. B. & Steven, A. C. (2013). J. Struct. Biol. 184, 226–236.

Chen, S., McMullan, G., Faruqi, A. R., Murshudov, G. N., Short, J. M., Scheres, S. H. W. & Henderson, R. (2013). Ultramicroscopy. 135, 24–35.

DiCiccio, C. J. & Romano, J. P. (2017). J. Am. Stat. Assoc. 112, 1211–1220.

Guo, H., Suzuki, T. & Rubinstein, J. L. (2019). Elife. 8, e43128.

Harauz, G. & Van Heel, M. (1986). Optik (Stuttg). 73, 146–156.

Heel, M. van & Schatz, M. (2017). BioRxiv.

Van Heel, M. & Schatz, M. (2005). J. Struct. Biol. 151, 250–262.

Hunter, J. D. (2007). Comput. Sci. Eng. 9, 90–95.

Juszkiewicz, S., Chandrasekaran, V., Lin, Z., Kraatz, S., Ramakrishnan, V. & Hegde, R. S. (2018). Mol. Cell. 72, 469–481.

Kucukelbir, A., Sigworth, F. J. & Tagare, H. D. (2014). Nat. Methods. 11, 63–65.

Kühlbrandt, W., Amunts, A., Liao, M., Cao, E., Julius, D., Cheng, Y., Allegretti, M., Li, X., Ban, N., Wimberly, B. T., Faruqi, A. R., Henderson, R., Daum, B., Walter, A., Horst, A., Osiewacz, H. D., Kühlbrandt, W., Schur, F. K. & Scheres, S. H. (2014). Science. 343, 1443–1444.

Lehmann, E. & Romano, J. (2005). Testing Statistical Hypotheses.

Massey, F. J. (1951). J. Am. Stat. Assoc. 46, 68–78.

Nieuwenhuizen, R. P. J., Lidke, K. A., Bates, M., Puig, D. L., Grünwald, D., Stallinga, S. & Rieger, B. (2013). Nat. Methods. 10, 557–562.

Oliphant, T. E. (2007). Comput. Sci. Eng. 9, 10–20.

Orlova, E. V., Dube, P., Harris, J. R., Beckman, E., Zemlin, F., Markl, J. & Van Heel, M. (1997). J. Mol. Biol. 271, 417–437.

Pettersen, E. F., Goddard, T. D., Huang, C. C., Couch, G. S., Greenblatt, D. M., Meng, E. C. & Ferrin, T. E. (2004). J. Comput. Chem. 25, 1605–1612.

Rosenthal, P. B. & Henderson, R. (2003). J. Mol. Biol. 333, 721–745.

Saxton, W. O. & Baumeister, W. (1982). J. Microsc. 127, 127–138.

Sindelar, C. V. & Grigorieff, N. (2012). J. Struct. Biol. 180, 26–38.

Spahn, C. M. T., Jan, E., Mulder, A., Grassucci, R. A., Sarnow, P. & Frank, J. (2004). Cell. 118, 465–475.

Vilas, J. L., Gómez-Blanco, J., Conesa, P., Melero, R., Miguel de la Rosa-Trevín, J., Otón, J., Cuenca, J., Marabini, R., Carazo, J. M., Vargas, J. & Sorzano, C. O. S. (2018). Structure. 26, 337–344.

Van Der Walt, S., Colbert, S. C. & Varoquaux, G. (2011). Comput. Sci. Eng. 13, 22–30.

Weis, F., Beckers, M., Hocht, I. & Sachse, C. (2019). EMBO Rep.

Zi Tan, Y., Baldwin, P. R., Davis, J. H., Williamson, J. R., Potter, C. S., Carragher, B. & Lyumkis, D. (2017). Nat. Methods. 14, 793–796.

Zivanov, J., Nakane, T., Forsberg, B. O., Kimanius, D., Hagen, W. J. H., Lindahl, E. & Scheres, S. H. W. (2018). Elife. e42166.

